# A resource for probing influences of perinatal factors on neurodevelopment in 3-10-years-old Chinese children

**DOI:** 10.1101/2023.09.28.560037

**Authors:** Yin-Shan Wang, Xue-Ting Su, Li Ke, Qing-Hua He, Da Chang, JingJing Nie, XinLi Luo, Fumei Chen, Jihong Xu, Cai Zhang, Shudong Zhang, Shuyue Zhang, Huiping An, Rui Guo, Suping Yue, Wen Duan, Shichao Jia, Sijia Yang, Yankun Yu, Yang Zhao, Yang Zhou, Li-Zhen Chen, Xue-Ru Fan, Peng Gao, Chenyu Lv, Ziyun Wu, Yunyan Zhao, Xi Quan, Feng Zhao, Yanchao Mu, Yu Yan, Wenchao Xu, Jie Liu, Lixia Xing, Xiaoqin Chen, Xiang Wu, Lanfeng Zhao, Zhijuan Huang, Yanzhou Ren, Hongyan Hao, Hui Li, Jing Wang, Qing Dong, Yanli Chen, Ruiwang Huang, Siman Liu, Yun Wang, Qi Dong, Xi-Nian Zuo

## Abstract

Adverse perinatal factors can disrupt the normal development of the brain, with potential long-term impacts on children’s overall development. Currently, the neuropathological mechanisms by which these factors lead to various neurodevelopmental disorders (NDDs) remain largely unknown. An open resource that integrate perinatal factors with brain and mental health development is essential for investigating NDD-related aetiology. In this Data Descriptor, we introduce a multicentre database containing information on perinatal factors that influence children’s brain-mind development, namely, periCBD, that combines neuroimaging and behavioural phenotypes with perinatal factors associated with a high incidence of NDDs at county/region/central district hospitals. PeriCBD was designed to establish a platform for the investigation of individual differences in brain-mind development among children aged 3–10 years are associated with perinatal factors. Ultimately, our goal was to develop an early prediction and screening model for NDDs that leverages normative data to facilitate NDD aetiology research. Herein, we provide a systematic overview of the data acquisition/cleaning/quality control/sharing, processes of periCBD and present preliminary brain-mind associations.

## Background & Summary

Neurodevelopmental disorders (NDDs) refer to a series of mental disorders that disrupt the motor and behaviour development of children in the kindergarten through twelfth grade (K-12 children)^1^. These disorders typically manifest in early childhood and impact various aspects of behavior, with no possibility of spontaneous recovery during development. Children with NDDs typically do not have physical disabilities, joint diseases, hormone disorders, or neurological disabilities. As a result, they often appear to exhibit typical development at the early stage of NDDs, leading to delays in intervention and treatment. NDDs may also influence the academic performance and mental health of children, resulting in abnormal performances that hinder typical development and cause considerable distress. In the absence of appropriate interventions, these problems may persist into adulthood. Advancements in neuroimaging technology have provided new opportunities to understand the neural mechanism and brain correlates of NDDs and how adverse perinatal factors lead to NDDs. It is therefore of utmost importance to establish a screening model that combines neuroimaging and behaviour phenotype data to facilitate early detection and intervention for NDDs.

The perinatal period refers to the time from 28 gestational weeks to one week after delivery. It is a critical phase in foetal and infant brain development characterized by rapid and complex development. Neurodevelopment during this period is crucial and lay the foundation for later cognitive, emotional and behavioural abilities. Various perinatal factors, including preterm birth, low birth weight, and other adverse conditions, are known to pose significant risks for NDDs. Previous studies have shown that children exposed to adverse perinatal factors are more susceptible to white matter dysplasia and have a higher likelihood of developing different types of NDDs^2–4^. For example, Davis and colleagues^5^ observed that children born very preterm (<28 gestational weeks) or with extremely low birth weight (ELBW <1000 g) had an approximately five times higher risk of developmental coordination disorder (DCD) than typically developing children at 8 years of age; and extremely preterm (<26 gestational weeks) children were 4.3 times more likely to develop attention-deficit/hyperactivity disorder (ADHD) than those born at term; moreover, extremely preterm children exhibited an increased risk of autism spectrum disorders (ASD)^6^. Despite these findings, shared and specific mechanisms by which perinatal factors mediate brain-behaviour relationships in different NDDs remain largely unknown.

To the best of our knowledge, there is currently no available open data resource designed to help study these mechanisms. To address this gap, we established a comprehensive multicentre brain-behaviour database called Perinatal Factors in Child Brain-mind Development (periCBD). This database encompasses a wide range of evaluations, including brain MRI scans, clinical manifestations of NDDs, and assessments of motor development. During the data collection process, we also considered other factors not associated with NDDs, such as intellectual, cognitive, or behavioural problems, to ensure that these factors were appropriately considered and controlled for. This helps us to better isolate and examine the specific influences of perinatal factors on child development. The investigation of perinatal factors, such as maternal age at delivery, duration of pregnancy, and birth weight, was carried out through the collection and analysis of clinical medical data as well as parental questionnaires. At the pilot stage, obstetric and paediatric data were collected from 2011 to 2018, recruitment was conducted at two district hospitals (the Anyang Maternal and Child Health Hospital of Henan Province and the People’s Hospital of Liangping District of Chongqing). Through the establishment of periCBD, we aimed to create a valuable resource that will contribute to a better understanding of the complex relationships among perinatal factors, brain development, and child behaviour. This database can serve as a foundation for future research and pave the way for advancements in the prevention, diagnosis, and treatment of NDDs.

## Methods

### Overall Design

The aim of periCBD is to build a representative, multidimensional brain-behaviour database comprising county-level data on Chinese children aged 3-10 years who experienced perinatal factors. The long-term goal of this project is to create a national resource that covers the 34 provinces in China, with 34 sites strategically distributed across counties and districts. In the initial phase, we selected two pilot sites: the Anyang Maternal and Child Health Hospital of Henan Province (AY site) and the People’s Hospital of Liangping District of Chongqing (LP site). The two sites were chosen to test the effectiveness of our recruitment and data collection methods. Our target population mainly included preschool children (3-6 years old) and school-age children (7-10 years old), as these age groups exhibit a higher incidence of NDDs that enables investigation of children with/without perinatal factors and collection of developmental data related to intelligence, motor skills and cognition. By integrating these developmental assessments with neuroimaging data, our aim is to provide a valuable resource for exploring the effects of perinatal factors on brain development. Additionally, we aimed to investigate the associations between of periantal factors with sensorimotor and cognitive development and the underlying mechanisms involved.

### Recruitment Strategy

Tow groups of participants were recruited for the study: the perinatal factor (PF) group and the normal control (NC) group. The PF group consisted of children aged 3-10 years who had experienced adverse perinatal factors, namely preterm birth (gestational age < 37 weeks) and/or low birth weight (birth weight < 2,500 g). To identify eligible individuals, obstetric and paediatric data from 2011 to 2018 were reviewed, and the eligibility of these children was further confirmed through telephone calls. The NC group comprised children without any adverse perinatal factors. They were recruited from local hospitals from June to August 2021. Parents were invited to attend seminars at the hospital, where they were provided with detailed information regarding the objectives and goals of the project. Additionally, they were informed about the scope of the project through posters that were displayed in various local communities. The NC children were selected to match the PF group in terms of age and demographic characteristics. During the recruitment process, exclusion criteria were applied to ensure that factors potentially affecting the child’s development, such as intellectual, psychiatric or behavioural problems that were not due to NDDs, were accounted for. All the data collected were later double-checked with the information provided by the caregivers of the individual participants to ensure accuracy. For a clearer understanding of the recruitment procedure, please refer to Figure 1.

**Figure 1.**
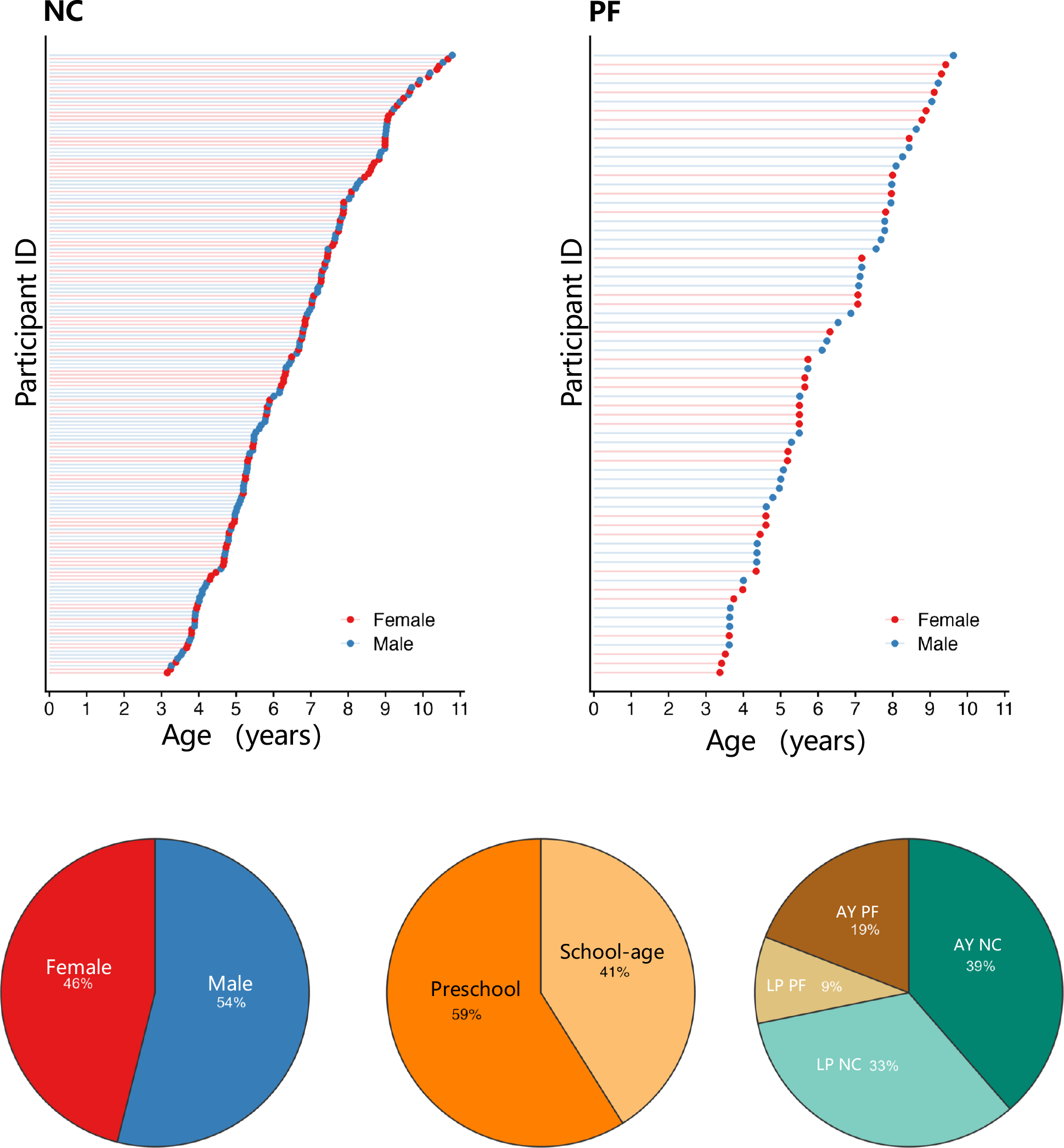
The demographic characteristics of individuals in the periCBD dataset. A total 241 children were involved in this study in the normal control (NC) group and the perinatal factor (PF) group. Among the participants, 46% were female and 59% were preschool children, while 19% of children in the AY group and 9% of children in the LP group had PF.

### Participants

A total of 241 children (age: 3-10 years; LP site: 74 preschool children, 41 school-age children; AY site: 68 preschool children, 58 school-age children) were recruited. There were 68 children in the PF group (AY site: 46; LP site: 22) and 173 children in the NC group (AY site: 80; LP site: 93). Further details about the demographic information are presented in Figure 1. All participants were enrolled in the study after obtaining consent from the children’s parents. This study was approved by the Institutional Review Board (IRB) of the State Key Laboratory of Cognitive Neuroscience and Learning at Beijing Normal University (file number: ICBIR_A_0206_001).

## Data acquisition

### Demographic, perinatal and family influences

The data collection process involved gathering information on various factors, including maternal age, body mass index (BMI), lifestyle, family socioeconomic status (SES), perinatal risk factors, maternal emotional problems, and parental behaviour. Maternal emotional problems were evaluated using multiple assessment tools, namely, the Edinburgh Postnatal Depression Scale^7^, the Self-Rating Depression Scale^8^, the Trait Depression Scale^9^, the Self-Rating Anxiety Scale^8^, and the State-Trait-Anxiety Inventory^10^. Parental behaviour was evaluated by the Life Event Scale (partial)^11^, Child Neglect Scale (partial)^12^, the Brief Coparenting Relationship Scale (Brief CRS)^13^, and Parenting Sense of Competence Scale (PSOC)^14^. For more details on the applicable age range and duration of these assessments/procedures, please refer to Table 1.

**Table 1.**
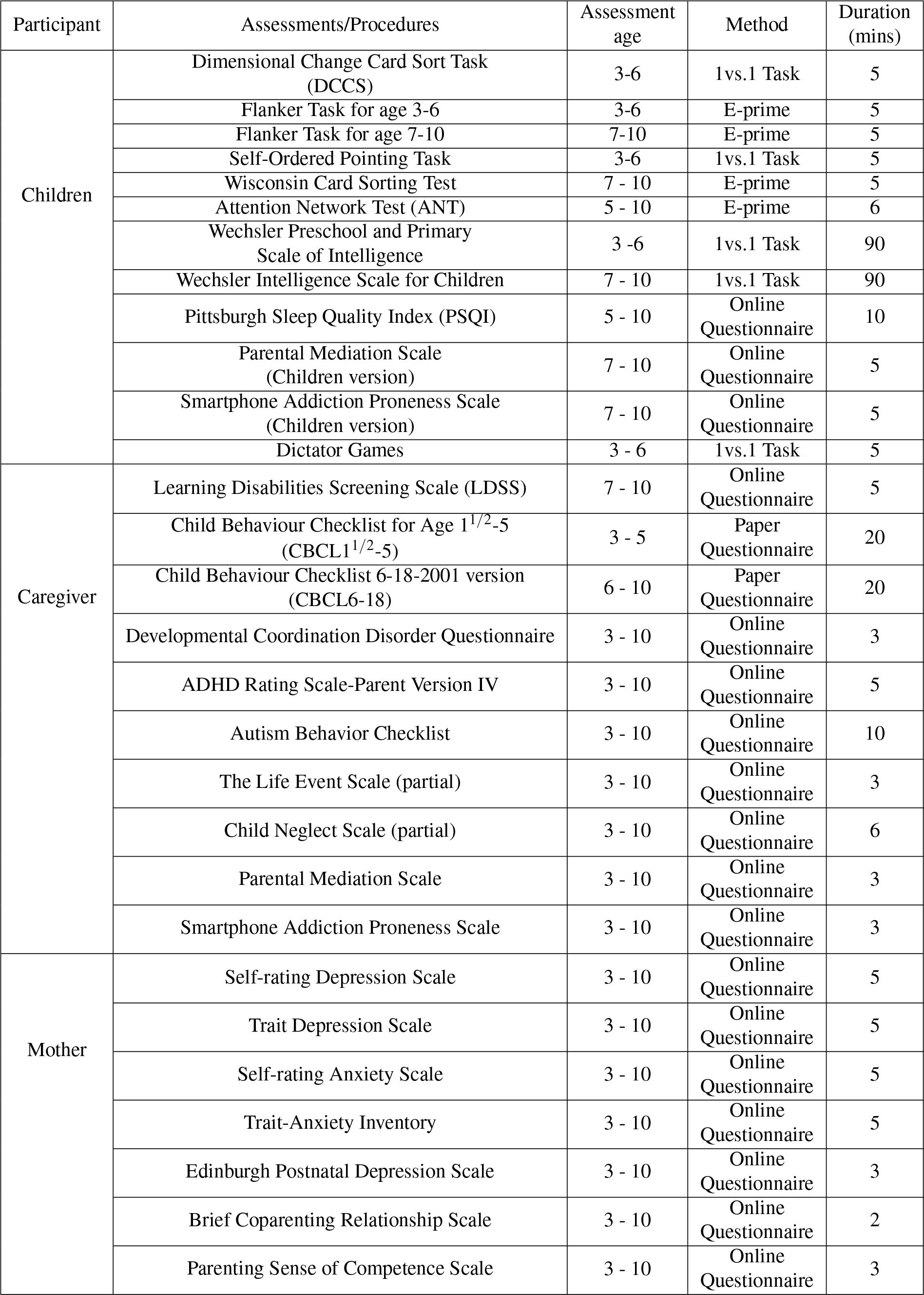
The duration infomation of Assessments/Procedures in periCBD.

- **Family Socioeconomic Status (SES)** Family socioeconomic status^15^ was assessed by synthesizing five items from three dimensions: parents’ education level, parents’ occupation, and family income. The caregiver provided the answers for this assessment. A higher score indicated a higher family SES. Children’s early family SES was measured using a four-item scale (e.g., The child grew up in a relatively wealthy neighbourhood)^16^. This scale effectively reflects the quality of the early developmental environment for children. The child’s mother was asked to recall the conditions during pregnancy and the first week of the child’s life. Scores ranged from 1 (strongly disagree) to 5 (strongly agree), with higher scores indicating a higher early family socioeconomic status.
- **Edinburgh Postnatal Depression Scale** The Edinburgh Postnatal Depression Scale is a self-report assessment tool specifically designed to evaluate postnatal depression in mothers. The scale is completed by the mother of the child and consists of 10 items. Each item is rated on a 4-point scale, ranging from 0 to 3. The total score ranges from 0 to 30, with higher scores indicating a higher severity of postnatal depression. The scale has a sensitivity of 86%, a specificity of 78%, a positive predictive value of 73%, and a split-half reliability of 0.88^7,17^.
- **Self-Rating Depression Scale and Trait Depression Scale** The Self-Rating Depression Scale and the Trait Depression Scale were administered to the child’s mother to assess her current depressive state and trait depression, respectively. The Self-Rating Depression Scale consists of 20 items, which are rated on a scale from 1 to 4. A higher total score indicates a more severe current depression symptoms in the mother^8^. The Trait Depression Scale is a 16-item scale, with items rated on a scale of 1 to 4, and total scores ranging from 16 to 64. A higher total score on this scale reflects a higher level of trait depression in the mother^10^. The two scales are widely used and have been proven to possess high validity in China.
- **Self-Rating Anxiety Scale and State-Trait-Anxiety Inventory** The Self-Rating Anxiety Scale and the State-Trait Anxiety Inventory were completed by the mother to assess her current state of anxiety and trait anxiety, respectively. The Self-Rating Anxiety Scale consists of 20 items, each rated on a scale of 1 to 4. A higher total score on this scale indicates more severe anxiety experienced by the mother^8^. The Trait Depression Scale is a 20-item scale with total scores ranging from 20 to 80. Higher scores indicate a higher level of trait anxiety in the mother^10^.
- **Parental behaviour** To assess various aspects of family dynamics and parental functioning, several scales were employed. First, the Life Event Scale (partial) was used to assess family stress. The caregiver answered 10 items, with higher scores indicating a greater amount of negative life events, i.e., greater perceived family stress. The Life Event Scale has good internal consistency reliability (0.90) and acceptable levels of construct validity (*χ*^2^*/d f* = 2.75, RMSEA=0.067, IFI=0.92, TLI=0.91, CFI=0.92)^11^ .The Child Neglect Scale (partial) measures the extent of neglect experienced by the child. It consists of 29 items organized into three subscales: physical neglect, affection neglect, and communication neglect. Caregivers provided responses for each subscale. Higher scores on each subscale indicate greater levels of neglect in the corresponding dimension. The scale was developed for the Chinese cultural context with an internal consistency reliability of 0.848, a split-half reliability of 0.810, and a test-retest reliability of 0.897. The construct validity is good (*χ*^2^*/d f* = 1.766, RMSEA=0.047, GFI=0.917, TLI=0.916)^12^.The Brief Coparenting Relationship Scale (Brief CRS) evaluates the quality of coparenting in families. It assesses the level of support and cooperation between fathers and mothers in parenting, to evaluate the quality of the family environment. The mother completes the scale, rating to 14 items on a scale rangingfrom 1 (not very true) to 5 (very true). Higher scores indicate higher levels of parental support and cooperation in coparenting. The scale has demonstrated good reliability and validity in Chinese studies^13^. Finally, the Parenting Sense of Competence Scale (PSOC) was used to assess maternal self-efficacy in parenting. It consists of 17 items with scores ranging from 1 (strongly disagree) to 6 (strongly agree). The scale is divided into a performance subscale and a satisfaction subscale. A higher overall score reflects a greater sense of efficacy in parenting. In China, the scale has exhibited good reliability, with internal consistency reliability values of 0.82, 0.80, and 0.85 for the total scale and efficacy and satisfaction subscales, respectively. Additionally, it has good construct validity (*χ*^2^*/d f* = 1.67, RMSEA=0.059, NFI=0.91, INNF=0.92, CFI=0.91, and IFI=0.94)^14^.

### Neurodevelopmental disorders (NDDs)

The periCBD dataset also collects information about neurodevelopmental disorders (NDDs) and incorporates multiple questionnaires to assess these conditions. The details and detection rates of these assessments can be found in Table 2.

**Table 2.**
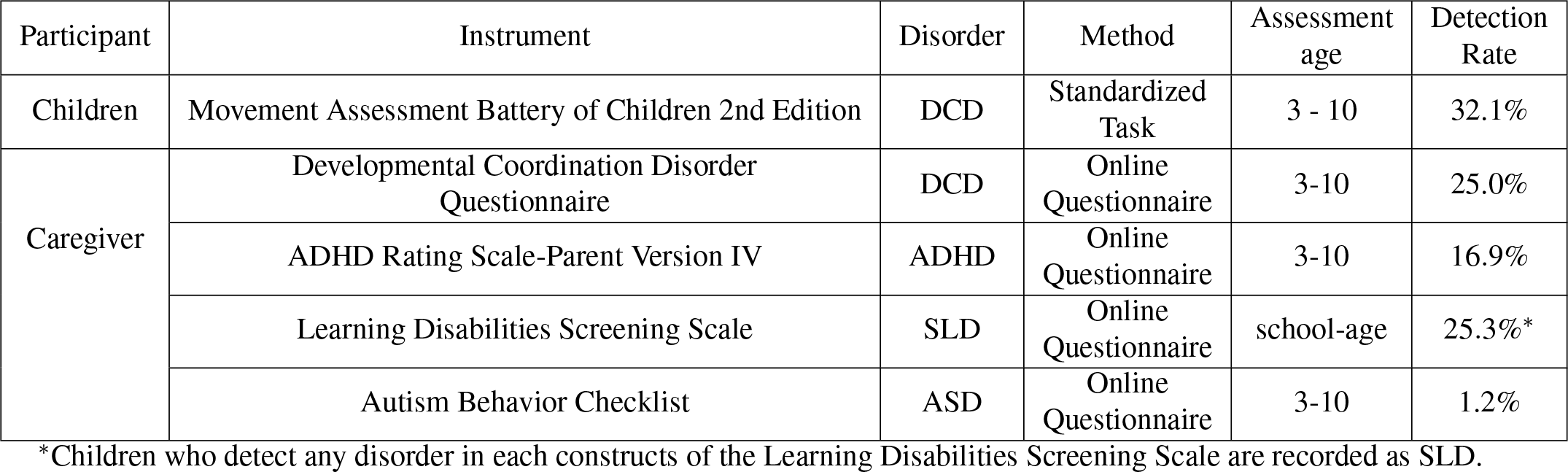
Detail information NDDs Assessments.

- **ADHD Rating Scale-Parent version IV (ADHD RS-IV)** The incidence of ADHD was assessed by ADHD Rating Scale-Parent version IV (ADHD RS-IV). The 18 ADHD RS-IV items are divided into the inattentive and hyperactive/impulsive dimensions. The results can be used to categorize individuals into four groups: no ADHD, ADHD predominantly inattentive type, ADHD predominantly hyperactive/impulsive type, and ADHD combined type. The Chinese version of the ADHD RS-IV has good internal consistency reliability, with a coefficient of 0.91. Additionally, it exhibits high construct validity (confirmatory factor analysis of the three-factor model: df =132, *χ*^2^ = 905, RMSEA=0.068, NFI=0.93, NNFI=0.91)^18,19^.
- **Autism Behavior Checklist (ABC)** The Autism Behavior Checklist (ABC) was used to evaluate whether the children had autism spectrum disorder. This questionnaire, consisting of 57 items, is completed by the caregiver and serves to evaluate various characteristics associated with autism. The ABC items are divided into five subscales: 1) sensory; 2) relating; 3) body and object use; 4) language; and 5) social and self-help.^20^.
- **Learning Disabilities Screening Scale (LDSS)** The Learning Disabilities Screening Scale (LDSS) was used to investigate the incidence of specific learning disorder (SLD) among the school-age children. The LDSS has 18 items divided into 6 constructs: reading aloud, reading comprehension, spelling, written expression, calculation, and mathematical reasoning. It is suitable for the assessment of learning disabilities in Chinese children and has exhibited good internal consistency and sufficient structural validity and criterion correlation validity^21^.
- **Assessment of motor development** The 2^*nd*^ Movement Assessment Battery for Children (MABC-2) and the Chinese version of the Developmental Coordination Disorder Questionnaire (DCDQ-C) were used to evaluate the motor development of children. The MABC-2 is a widely-used standardized assessment tool for measuring children’s motor development. It is suitable for use in children aged 3-16 years and has three dimensions: manual dexterity, aiming and catching, and balance. Ke Li^22^established the norm for urban children in China, reporting an internal consistency reliability coefficient of 0.583, which is close to the acceptable range. CFI and TFI values of the scale are both approximately 0.9, and RMSEA is 0.045, indicating good structural validity^23^. The DCDQ-C, completed by the caregiver, is divided into three dimensions: control during movement, fine motor/handwriting, and general coordination. Zhu Qingqing and colleague^24^ applied it in Chinese children aged 4-6 years and found an internal consistency reliability coefficient of 0.867, indicating good reliability. The questionnaire also has good structural validity^25^.

### Behavioural problems

- **Child Behaviour Checklist** Children’s behavioural problems were assessed by the Child Behaviour Checklist 6-18-2001 version (CBCL6-18) and Child Behaviour Checklist for Age 1^1*/*2^*−* 5(CBCL1^1*/*2^ *−* 5)^26,27^. These questionnaires were completed by the caregiver.
- **Pittsburgh Sleep Quality Index** The Pittsburgh Sleep Quality Index was used to measure the participant’s sleep quality (in children 5-10 years old), A higher total score indicates poorer sleep quality^28^.
- **Parental Mediation Scale** The Parental Mediation Scale was adapted from the Television Mediation Scale, and was completed by the caregiver and by children aged 7-10 years. It consists of 7 items, with higher total scores indicating increased parental monitoring. The original Television Mediation Scale comprises 15 items and is divided into three dimensions: instructive mediation, restrictive mediation, and social coviewing. However, for the purpose of this project, the scale was simplified, focusing mainly on restrictive mediation and social coviewing^29^.
- **Smartphone Addiction Proneness Scale** Caregivers and children aged 7-10 years completed the Smartphone Addiction Proneness Scale. This 15-item scale measures smartphone addiction across four subdomains: disturbance of adaptive functions, virtual life orientation, withdrawal, response, and tolerance. The scale has demonstrated good reliability within primary and secondary school populations^30^. Additionally, it has been found to have good reliability specifically within Chinese children and adolescents^31^.
- **Dictator Game** The dictator game was used to assess the sharing behaviour of children aged 3-6 years. This activity involves providing children with 30 stickers and requesting them to select their favourite 10 stickers. Once they have made their selection, they are informed that they have the option to give some of these 10 stickers to another child who will join later. It is emphasized to the children that they are not obligated to give away their stickers. They are then instructed to place the stickers they choose to keep for themselves in a brown envelope and the stickers they decide to give to others in a white envelope. The number of stickers given to others is recorded for analysis^32^.

### Intelligence assessment

The Wechsler Preschool and Primary Scale of Intelligence-fourth edition (*WPPSI − IV*) and the Wechsler Intelligence Scale for Children-fourth edition (*WISC− IV*) were used to measure the intelligence of children aged 3-6 and 7-10 years, respectively^33^. The Chinese version of the WPPSI-IV, revised in 2014, is applicable to children aged between 2 years and 6 months and those aged 6 years and 11 months. The WPPSI-IV provides composite scores for full-scale IQ (*FSIQ*) and on 5 indices: the verbal comprehension index (*VCI*), visual spatial index (*VSI*) working memory index (*WMI*), fluid reasoning index (*FRI*), and processing speed index (PSI). In this study, measures of verbal comprehension, visuo spatial skills and working memory were administered to children aged 3 years and 0 months to 3 years and 11 months. For children aged 4 years and 0 months to 6 years and 11 months, measures of fluid reasoning and processing speed were also included. The Chinese version of *WISC − IV*, revised in 2007, is suitable for children aged between 6 years and 0 months and 16 years and 11 months. The composite scores obtained from the WISC-IV include full-scale IQ (*FSIQ*) and 4 indice: the verbal comprehension index (*VCI*), perceptual reasoning index (*PRI*), working memory index (*WMI*) and processing speed index (*PSI*). The four index scales of the *WISC IV* were utilized to measure the intelligence of children aged 7 years and 0 months to 10 years and 11 months in this study.

### Cognitive Tasks/Tests

A battery comprising eight tasks was employed to evaluate children’s cognitive ability.

- **Dimensional Change Card Sort Task** This task was used to assess the cognitive flexibility of children aged 3 to 6 years. In a standard DCCS task^34^, children are instructed to first sort a set of cards (e.g., red rabbits and blue boats) in Phase 1 based on a preconversion rule. They are then instructed to resort the same set of cards in Phase 2 based on a postconversion rule. In Phase 3, participants are presented with a new set of cards (e.g., green flowers and yellow cars) and are instructed to classify them based on the rules established in Phase 2. In Phase 4, they are again presented with new cards and asked to classify them based on the rules from Phase 1. For example, children may be instructed to initially sort the cards based on colour, followed by sorting them based on shape. Subsequently, they are instructed to sort new cards by shape again, and finally, by colour once more. The number of correctly classified cards are counted at each stage. A higher score indicates better cognitive flexibility in children. The test materials were presented using PowerPoint, and children simply pointed to the selected cards on the screen with their hands.
- **Flanker Task** This task was employed to investigate children’s inhibitory control ability^35^. In the task, a white fixation cross “+” in the centre of a black screen for 1000 ms. After a 500 ms interval, a white target stimulus was presented. The specific target stimulus varied depending on the age of the children (fish for children under 7 years old, arrows for children 7 years old and older). The target stimulus consisted of five arrows (or fish), and the orientation of the central arrow (fish) either matched or did not match the orientation of the arrows (or fish) on either side. The children were instructed to focus on the direction of the central arrow (or fish) while ignoring the direction of the arrows (fish) on both sides. They were then instructed to respond quickly and accurately by pressing the “F” key if the direction was to the left or the “J” key if the direction was to the right. The inhibitory control ability of the participants was measured as the difference in response time between the inconsistent condition and the consistent condition. A greater difference in response time indicated poorer inhibitory control ability.
- **Wisconsin Card Sorting Test** The Wisconsin Card Sorting Test was implemented to measure cognitive flexibility in children aged 7-10 years^36^. During the test, children were presented with a computer screen that displayed four templates at the top: 1 red triangle, 2 green pentagrams, 3 yellow crosses, and 4 blue circles. A response card with varying shapes (triangle, pentagram, cross, circle), colours (red, yellow green, blue), and numbers (1, 2, 3, 4) would then appear randomly in the centre of the screen. Children were instructed to classify the response cards based on the four stimulus card templates. The test did not explicitly reveal the classification rule to the children; instead, they were informed whether each choice was correct or incorrect. The WCST yields several evaluation indicators: (1) total number of responses, (2) number of completed classifications, (3) number of incorrect responses, and (4) number of responses required to complete the first classification. A lower total number of responses, a greater number of completed classifications, fewer number of incorrect responses, and a smaller number of responses required to complete the first classification indicated better cognitive flexibility in children.
- **Self-Ordered Pointing Task** The task was used to measure working memory span in children aged 3-6 years. In this task, children were presented with a book containing pictures. Initially, they viewed a page with two pictures and selected one of them. Subsequently, the researchers turned to the next page, which displayed the same two pictures as before but in different positions. The children were then instructed to point to the picture they had not previously chosen. This process was repeated with each new page introducing a new image placed alongside the original two, which were arranged differently from the previous page. The children had to identify the image that had not been selected in each arrangement. This continued until the child made an incorrect selection twice in a row after the images were rearranged. At that point, the task was concluded, and the number of images presented before task termination determined the working memory span of the participant.
- **Attention Network Test(ANT)** The classic attention network test (ANT)^37^ was applied to assess the three attention networks: alerting, orienting and executive attention. Each trial commenced with a “+” symbol displayed at the centre of the screen, serving as a reminder for children to maintain their focus on the centre of the screen throughout the experiment. Subsequently, an asterisk “*” appeared as a cue in the central position. There were four types of cues: no clue, central clue, double clue (two clues appeared above and below the fixation point at the same time) and spatial clue (one clue appeared above or below the fixation point). Instead of arrows, images of cartoon fish were employed to indicate a direction (left or right) according to the direction in which the head of the centre “fish” pointed, as a preference for children. The children were instructed to determine the direction of the centre “fish’s” head (left or right) as quickly and accurately as possible. Response times (RTs) and accuracy were recorded for each trial. The effect value of the alerting network was computed by subtracting the response time in the two-cue condition from the response time in the no-cue condition. The effect value of the orienting network was calculated as the difference response time in the spatial cue context and the central cue context. The effect value of the executive control network was determined by subtracting the response time under consistent conditions from the response time under inconsistent conditions.
- **Intertemporal choice task** This task was designed to measure the intertemporal decision-making ability in children. During the task, children were presented with multiple choices in which they had no indicate their preference for receiving a smaller immediate reward (2, 4, or 6 gold coins) or a larger fixed reward (8 gold coins) after varying delays (5, 15, 25 or 40 seconds). For example, in some trials, the children had to decide between obtaining 6 gold coins instantly or waiting for 25 seconds to receive 8 gold coins. All participants underwent testing in the same pseudorandom order. Each immediate small reward was paired twice with each delayed large reward, resulting in a total of 24 choice trials. The options were visually represented by two planes displayed on a computer screen, each indicating a specific number of gold coins. The delay was conveyed through the altitude at which the aircraft was flying, where a higher altitude indicated a longer delay. The positioning of the delayed reward aircraft on the left and right sides was balanced throughout the experiment. Participants made their choice by pressing a corresponding button (right or left), and the chosen gold coins were then “dropped” into a basket at the bottom of the screen either immediately or after the specified delay. After each trial, the computer visually updated the total number of coins accumulated before proceeding to the next trial. The participants were informed about the total number of trials they would complete, but they were not explicitly informed about the duration of the delays. Instead, they learned the length of each type of delay during the practice task, allowing them to mentally associate each aircraft altitude with the corresponding delay time. Upon completion of the task, participants received the gold coins they had accumulated. The intertemporal decision-making ability was evaluated by calculating the area under the curve (AUC), which determined the “indifference point” for each delay time. A larger AUC indicated a greater willingness to wait and a higher level of intertemporal decision-making in children.
- **Children’s Gambling Task** Kerr and Zelazot^38^ adapted the Iowa gambling task to create a simplified gaming task for measuring hot executive function in children. This research method is considered complex but effective. The task involving two decks of cards, one with vertical bars and the other with polka dots on the front. Turning the cards over reveals a happy face or a sad face on the back. However, there is a difference between the decks. The cards with vertical bars mostly have a happy face on the back and occasionally have a sad face, while the cards with polka dots mostly have two happy faces on the back but sometimes have four, five, or six sad faces. In this task, a happy face representes win, and the number of happy faces represent the number of coins won. A sad face represents a loss, and the number of sad faces represent the number of coins lost. Coins could be exchanged for a candy. Participants can choose only one card per trial. The cards with vertical bars yield fewer coins each time (only one) but also lead to fewer coins lost on average (only one). In contrast, the cards with polka dots yield more coins each time (two) but also lead to many more on average (losses of four, five, or six). Therefore, choosing the cards with vertical bars is advantageous in the long run, while choosing the cards with polka dots is disadvantageous. In the experiment, the children are told to try to win as many coins as possible by the end of the “game” (which consistes of 50 card picks, unknown to the children beforehand). The first 25 choices are interpreted as children’s initial experience with the two types of cards, while the next 25 choices are used to assess emotional decision-making. The key dependent variable in this experiment was the affective decision index, which was calculated by subtracting the number of unfavourable cards chosen from the number of favorable cards chosen in trials 26 to 50. A higher affective decision index indicates stronger affective decision-making abilities in the participant.
- **Cake Gambling Task** This task was used to measure children’s risky decision-making abilities and is one of the research methods used to evaluate hot executive function^39^. The task involves dividing a cake into six pieces, with four pieces of one flavour (chocolate or strawberry) and two pieces of the other flavour. On the left and right sides of the cake, there are squares of pink and brown displaying different amounts of gold coins. The children are instructed to press a button to choose which flavour to bet on. If they make the correct choice, they receive a reward, but if they lose, they receive nothing. Throughout the gambling process, the cake with more pieces (4 pieces) is considered the low-risk option, while the cake with fewer pieces (2 pieces) is considered the high-risk option. The reward for the high-risk option (token =2/4/6/8) is always twice that for the low-risk option. Therefore, the expected value (EV), calculated as the probability multiplied by the reward magnitude, is equal for both options. The total risky decision index was determined by calculating the proportion of the risky option. A higher value indicates a greater inclination towards risk decision-making and poorer risky decision-making abilities.

### Magnetic resonance imaging (MRI)

All paediatric MRI scans were conducted at two different hospitals: the Peoples Hospital of Liangping District, Chongqing, using a 1.5T GE SIGNA_Creator scanner, and the Henan Anyang Maternal and Child Health Care Hospital, using a 3.0T GE SIGNA_Pioneer scanner. Details about the imaging protocols are presented in Table 3. During the scanning process, participants were instructed to remain still and not to fall asleep. Two 3D morphometric sequences (T1 and T2), a resting-state fMRI sequence, a diffusion tensor imaging (DTI) sequence and an arterial spin labelling (ASL) sequence were leveraged to collect the MRI data. The total scan time for each session was less than 30 minutes.

**Table 3.** MRI Protocols.

| Site | Sequence | TR (s) | TE (s) | FOV (mm) | FA (°) | Slice Thickness (mm) | Slice Spacing (mm) |
| --- | --- | --- | --- | --- | --- | --- | --- |
| AY | 3D T1 | 0.006736 | 0.002676 | 256 | 12 | 1 | 1 |
|  | 3D T2 | 3.202 | 0.092036 | 256 | 90 | 1 | 1 |
|  | BOLD | 2 | 0.03 | 224 | 90 | 3.9 | 4.5 |
|  | DTI | 12.8 | 0.078 | 224 | 90 | 2 | 2 |
|  | ASL | 4.711 | 0.11 | 224 | 111 | 3.5 | 3.5 |
| LP | 3D T1 | 0.006736 | 0.002676 | 256 | 12 | 1 | 1 |
|  | 3D T2 | 3.202 | 0.092036 | 256 | 90 | 1 | 1 |
|  | BOLD | 2 | 0.03 | 224 | 90 | 3.9 | 4.5 |
|  | DTI | 12.8 | 0.078 | 221 | 90 | 2 | 2 |
|  | ASL | 4.711 | 0.11 | 224 | 111 | 3.5 | 3.5 |

- **Morphometric MRI** Morphometric imaging consisted of a 3D T1 and a 3D T2 scan at both the AY and LP sites. The T1 images were collected with 1 mm isotropic voxels. The T1 sequence lasted for 4 minutes and 5 seconds in the SIGNA_Pioneer scanner, while it took 6 minutes and 04 seconds in the SIGNA_Creator scanner. For the 3D T2 image, a slice thickness of 1 mm was used. The acquisition time for this sequence was 4 minutes and 36 seconds on the SIGNA_Pioneer scanner, and 5 minutes and 59 seconds on the SIGNA_Creator scanner.
- **Resting-state fMRI** During the resting state protocol, participants were instructed to keep their eyes closed, remain motionless, and refrain from thinking about anything in particular. Resting state sequences with an 8-minute duration were acquired using a 3.9 mm resolution and a 0.4 mm gap at the AY site and a 3.5 mm resolution and 0.4 mm gap at the LP site.
- **Structural DTI** DTI scans on the 1.5T SIGNA Creator scanner consisted of 64 diffusion directions with a b-value equal to 1000 s/mm2 and 2 b-0 images, with 2.3 mm isotropic voxels. The acquisition time of data collection in the AP axis was 5 minutes and 43 seconds. DTI scans on the 3.0T Pioneer scanner consisted of 64 diffusion directions with a b-value equal to 1500 s/mm and 2 b-0 images, with 2 mm isotropic voxels, data collected in the AP direction. The aquisition time was 9 minutes and 3 seconds.
- **ASL MRI** The final sequence in each session was a pseudocontinuous arterial spin labelling (pCASL) sequence. Participants were asked to keep their eyes closed and remain motionless during the scan. The scank took 4 minutes and 14 seconds in the AY site, with a 3.5 mm resolution, and took 4 minutes and 32 seconds in the LP site, with a 3.5 mm resolution.

## Data Records

### Data Organization

In this dataset, the data file for each participant consisted of various imaging modalities, including structural imaging (T1 and T2 weighted images), resting-state functional MRI (rs-fMRI), diffusion tensor imaging and arterial spin labelling (ASL) perfusion imaging. The original data for each participant was converted into the NIFTI format and organized following the brain imaging data structure (BIDS) standard^40^. To ensure privacy, the facial information in each structural MRI was anonymized using the FaceMasking kit^41^, which obscures or removes identifiable facial features from the images.

### Quality Assessment

#### Phenotypic Data

As shown in Figure 2, all children were well clustered into the PF and NC groups based on their gestational age and birth weight. All the psychological and behavioural data are made available to database users, regardless of data quality. In addition, we have provided quality control (QC) files for each behavioural task and questionnaire, enabling users to make informed decisions about including specific data in their studies. The distributions of scores on the intelligence scale and the CBCL (both the social ability score and total score) are summarized in Figure 2.

**Figure 2.**
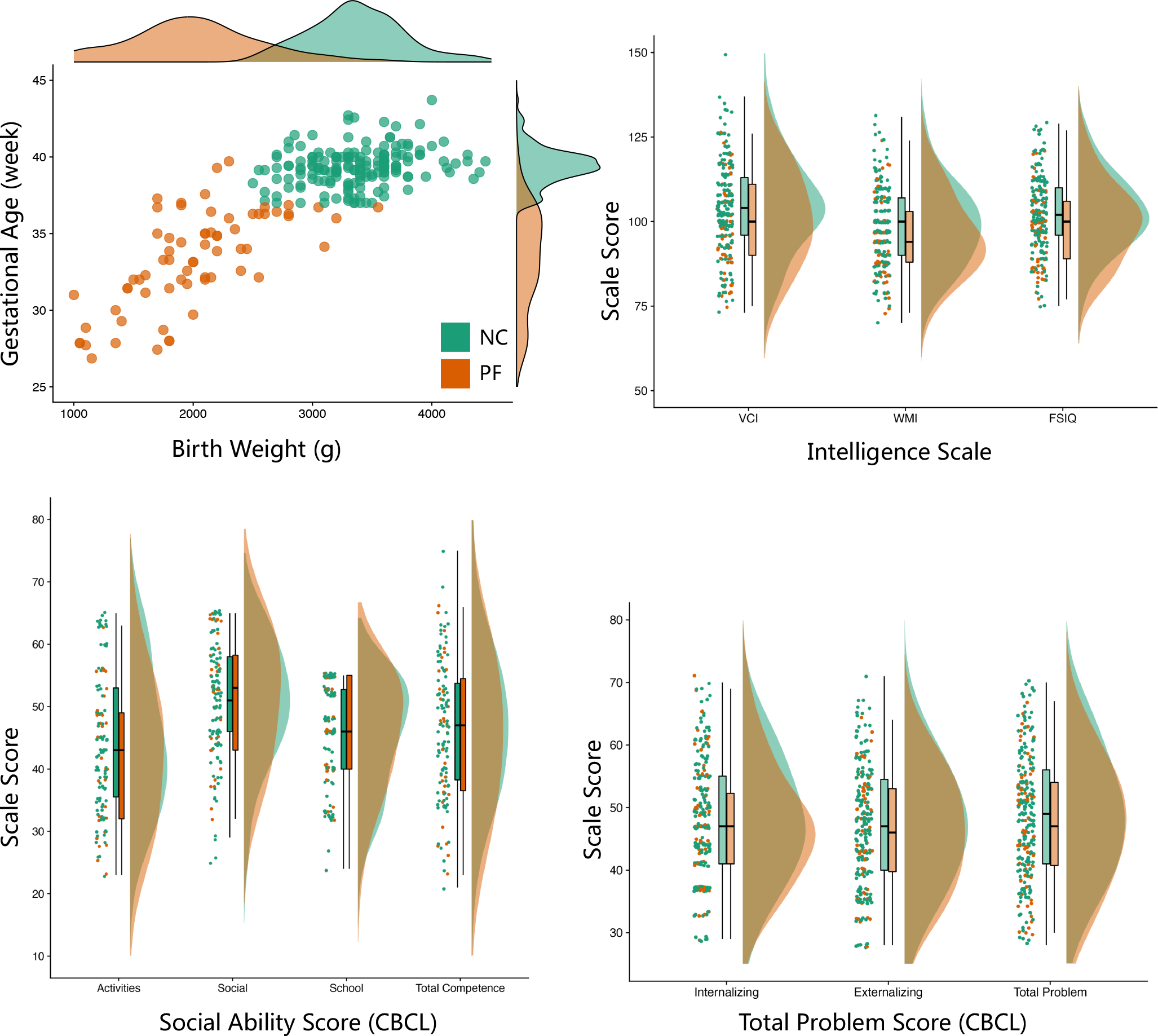
Distributions of non-neuroimaging measures. As depicted in the top-left panel, perinatal factors, including birth weight and gestational age differentiated children with perinatal factors (PF) from normal controls (NC). General intellectual ability (IQ) is visualized as (top -right) its standard score, including the verbal comprehension index (VCI), working memory index (WMI) and full-scale IQ (FSIQ). Mental health was evaluated by the Children Behaviour Check List (CBCL) in terms of social competence(bottom -left), including activities, social, and school competence, as well as total problems (bottom -right), including internalizing and externalizing problems.

**Figure 3.**
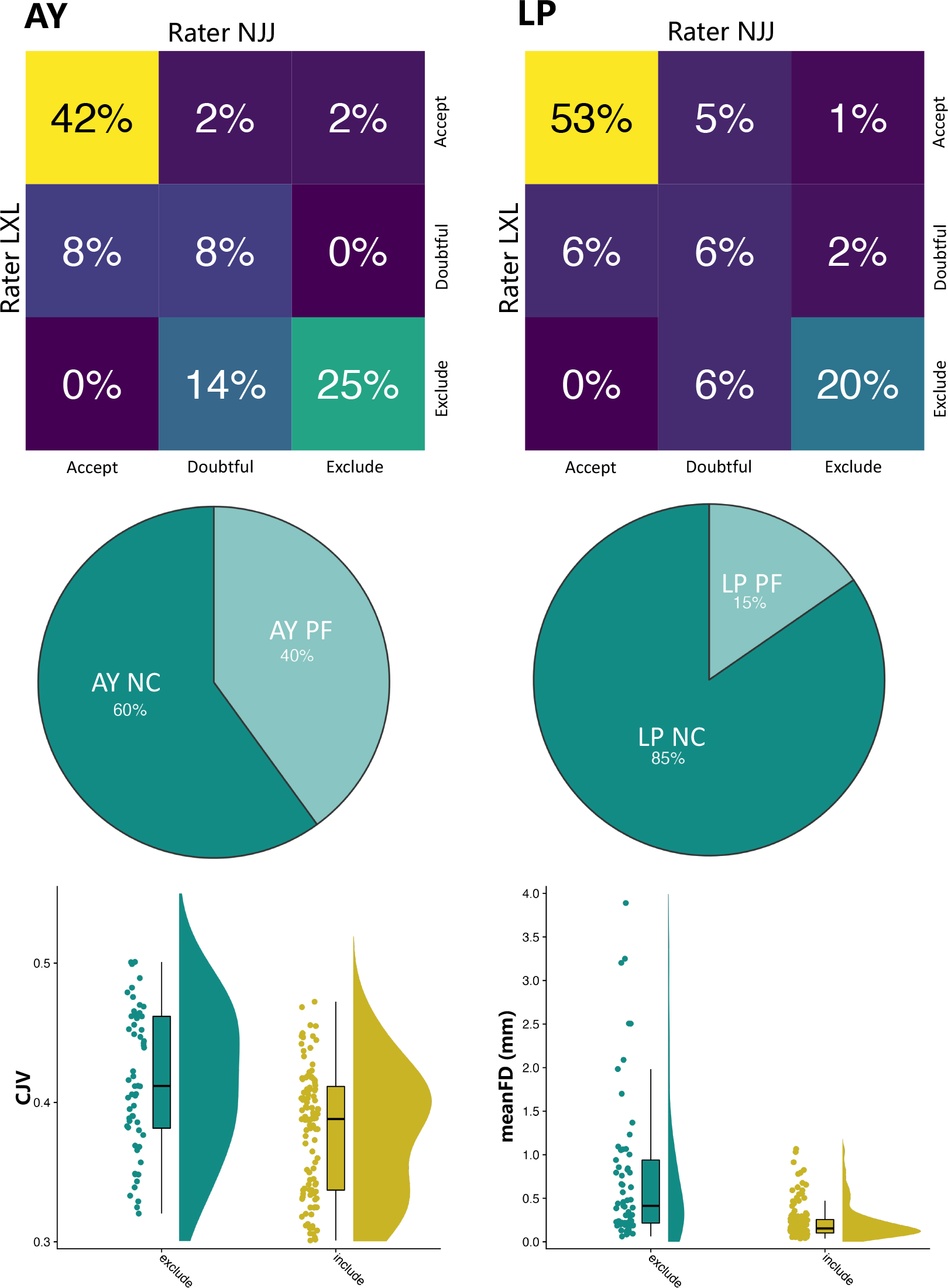
Summary of automatic and manual quality control of the images. The top panel shows heatmaps of the overlap percent of quality labels assigned by rater NJJ and rater LXL for images from the AY and LP sites. The middle panel shows pie charts of the percentages of NC and PF children whose images were excluded. The bottom panel indicates significant differences in both the structural coefficient of joint variation (CJV) and mean functional framewise displacement (meanFD) between the excluded images and included images.

#### MRI Datasets

Among the 241 children, 184 (89 at the AY site, 95 at the LP site) underwent an MRI scan and were subsequently included in the quality control procedure. The MRI Quality Control tool (MRIQC), an automatic quality control pipeline, was utilized to obtain image quality metrics (IQMs), such as the signal-to-noise ratio (SNR), coefficient of joint variation (CJV), mean framewise displacement (meanFD), and temporal SNR (tSNR)^42^. Similar to previous studies^4342^, manual rating procedures were conducted by two independent raters (coauthors: NJJ and LXL) who assessed all T1w images using an ordinal scale ranging from 0 to 2. They were aided by individual T1w reports generated from the MRIQC. In this rating framework, a rating of “0” indicated that the images were unusable and contained substantial artefacts. On the other hand, a rating of “2” indicated that the images were free from visible artefacts. Images with a rating of “1” had some artefacts present but were still considered usable.

### Data acquisition

The neuroimaging data that successfully met the quality control criteria are accessible through Chinese colour Nest Project (CCNP) – Lifespan Brain-Mind Development data Community under the title “perinatal factors in child brain-mind development, periCBD” (https://doi.org/10.57760/sciencedb.10690). The corresponding behaviour data have been published and can be found at the following DOI: (https://doi.org/10.57760/sciencedb.j00001.00423). For organizational purposes, all behaviour data are stored independently according to the participant ID. Each participant’s data comprise 10 files containing information on a total of 208 variables. These variables encompass a wide range of information, including demographics characteristics, cognitive abilities, intelligence measures, motor development assessments, disorder detection rates, partental behaviour, behavioural patterns, perinatal risk factors, and maternal emotional disorders.

## Technical Validation

### Structural Image Preprocessing

After two raters independently scored the images, the T1w images were excluded if at least one rater provided the “0” label. Figure 6 presents the proportion of the quality labels assigned by the two raters. The T1w images that passed QC were preprocessed using the Connectome Computation System (CCS)^44^ in the following steps: 1) Images were cropped with FSL’s **robustfov** to remove lower head and neck; 2) A spatially adaptive non-local means method was applied to denoise the images; 3) Skull stripping was performed using *deepbet*,^45^ which involved training a new pediatric U-Net model based on the CCNP dataset^46^; and 4) FreeSurfer (version 6.0.0) was utilized to obtain morphological measurements of different brain morphometry^47^. During this step, the brainmask generated by FreeSurfer was replaced by the brainmask generated by *deepbet*.

### Functional Image Preprocessing

All resting-state fMRI data were preprocessed by CCS. The specific details of the data preprocessing are beyond the scope of this study. In brief, the rs-fMRI pipeline consisted of the following steps: (1) dropping the first 10*s* (5 TRs) for the equilibrium of the magnetic field; (2) correcting head motion by FSL’s *mcflirt*; (3) slice timing correction; (4) de-spiking; (5) aligning functional images to high resolution T1 images using boundary-based registration (FreeSurfer’s *bbregister*); (6) mitigating nuisance effects such as ICA-AROMA-derived, CSF and white matter signals^48^; (7) removing linear and quadratic trends of the time series; (8) projecting volumetric time series to *fsaverage5* cortical surface space.

### Quality Assessment

To assess the level of agreement between the raters, we utilized the percentage of agreement method, given by:

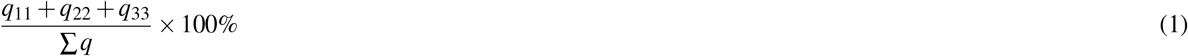

This equation calculates the sum of the diagonal element of the rating matrix *Q* and divides it by the sum of all elements of the rating matrix shown in Figure 2. This percentage indicates the extent of agreement between the raters. In addition, weighted cohen’s kappa values (*κ*) were also calculated to further evaluate inter-rater agreement in terms of the initial visual ratings. The obtained results demonstrated a high degree of consistency between raters in evaluating T1 images at both sites. The inter-rater agreement was found to be 74.1% for the AY site and 79.6% for the LP site. Furthermore, the weighted Cohen’s kappa value was 0.796, with a 95% confidence interval ranging from 0.702 to 0.889 at the AY site. Similarly, at the LP site, the weighted Cohen’s kappa value was 0.838, with a confidence interval ranging from 0.758 to 0.919.

Based on the manual ratings, a total of 35 out of 89 images were excluded at the AY site and 26 out of 95 images were excluded at the LP site. Among the excluded images, 40% belonged to PF participants at the AY site, and 15% were from PF participants at the LP site (Figure 2, middle row). Chi-square (*χ*^2^) goodness of fit tests were conducted to test whether the frequencies of images from PF and NC participants excluded via QC measures were significantly different from the frequencies in the original dataset. The results indicated that manual rating did not lead to a greater proportion of PF images excluded(AY site, *p* = 0.083, LP site, *p* = 0.48).

To investigate differences between the images labelled unusable and the included images in terms of head motion, we performed two-sample t tests on the coefficient of joint variation (CJV) of structural images and the average framewise displacement (meanFD) of resting-state images. More head motion was found in the excluded images (Figure 2, bottom row). The excluded images exhibited significantly higher CJV (*t* = 6.501, *p* = 7.44 *×* 10^*−*10^) and meanFD values(*t* = 5.233, *p* = 4.57 *×* 10^*−*7^) than the included images.

To further examine potential differences in image quality between the PF and NC groups after images passed QC, we performed two-sample t tests on several IQMs derived from MRIQC (structural: CJV, CNR, EFC, FBER, SNR_total, rs-fMRI: DVARS_std,FD_mean,tSNR,GSX_x,GSR_y). More details on the definition of IQMs can be found at the website of MRIQC (https://mriqc.readthedocs.io/en/latest/measures.html). We performed separate analyses for the AY and LP sites since the data were collected from two different sites with distinct scanners, field strengths, and protocols. Across all 10 IQMs, there were no significant differences observed between the PF and NC groups at both the AY and LP sites (Figure 4).

**Figure 4.**
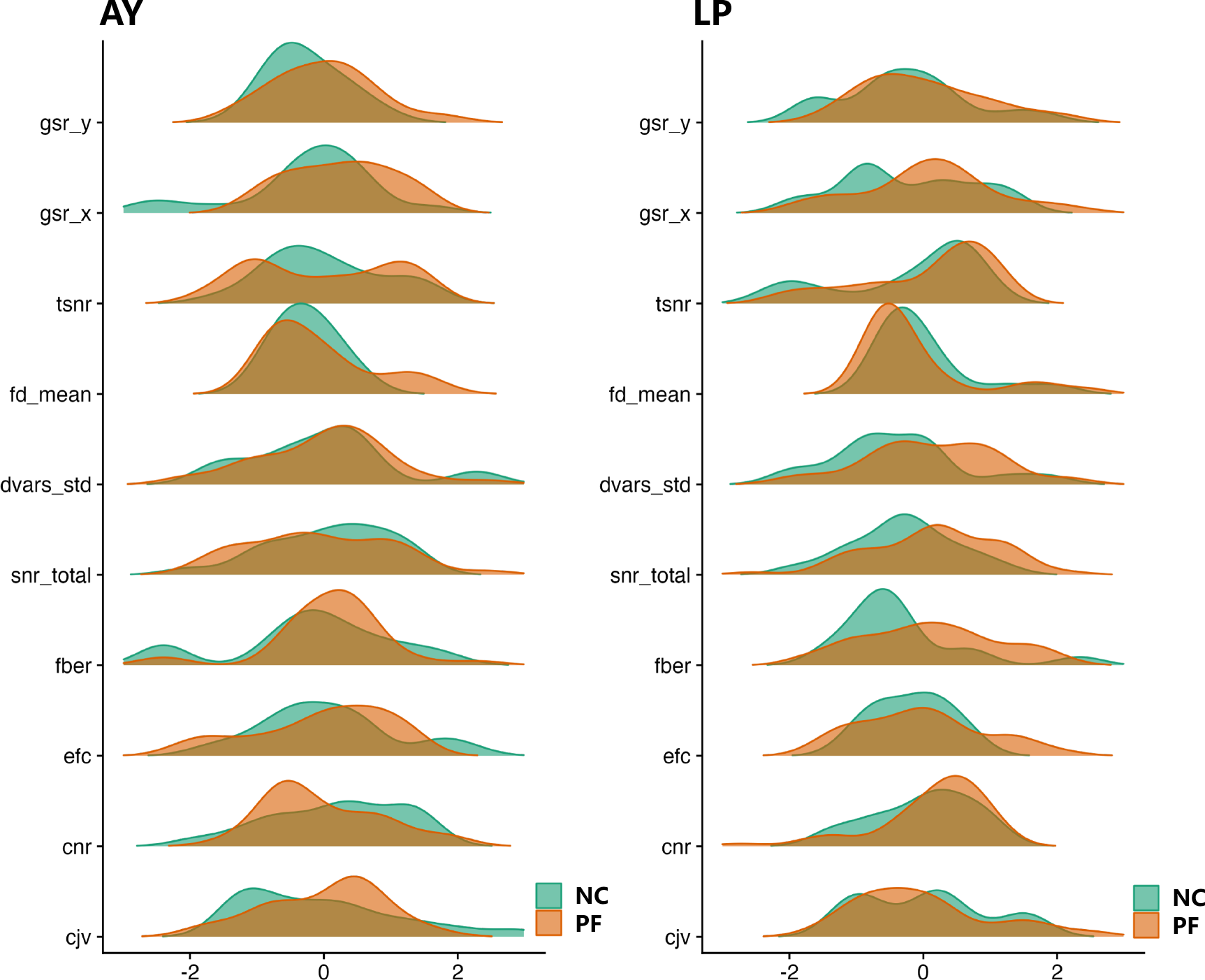
The distribution of the ten image quality metrics (IQMs) after z-transformation for images of children in the PF and NC groups at both th AY and LP sites that passed visual quality control. Two-sample t tests revealed no significant differences in any of the IQMs between the NC and PF groups.

**Figure 5.**
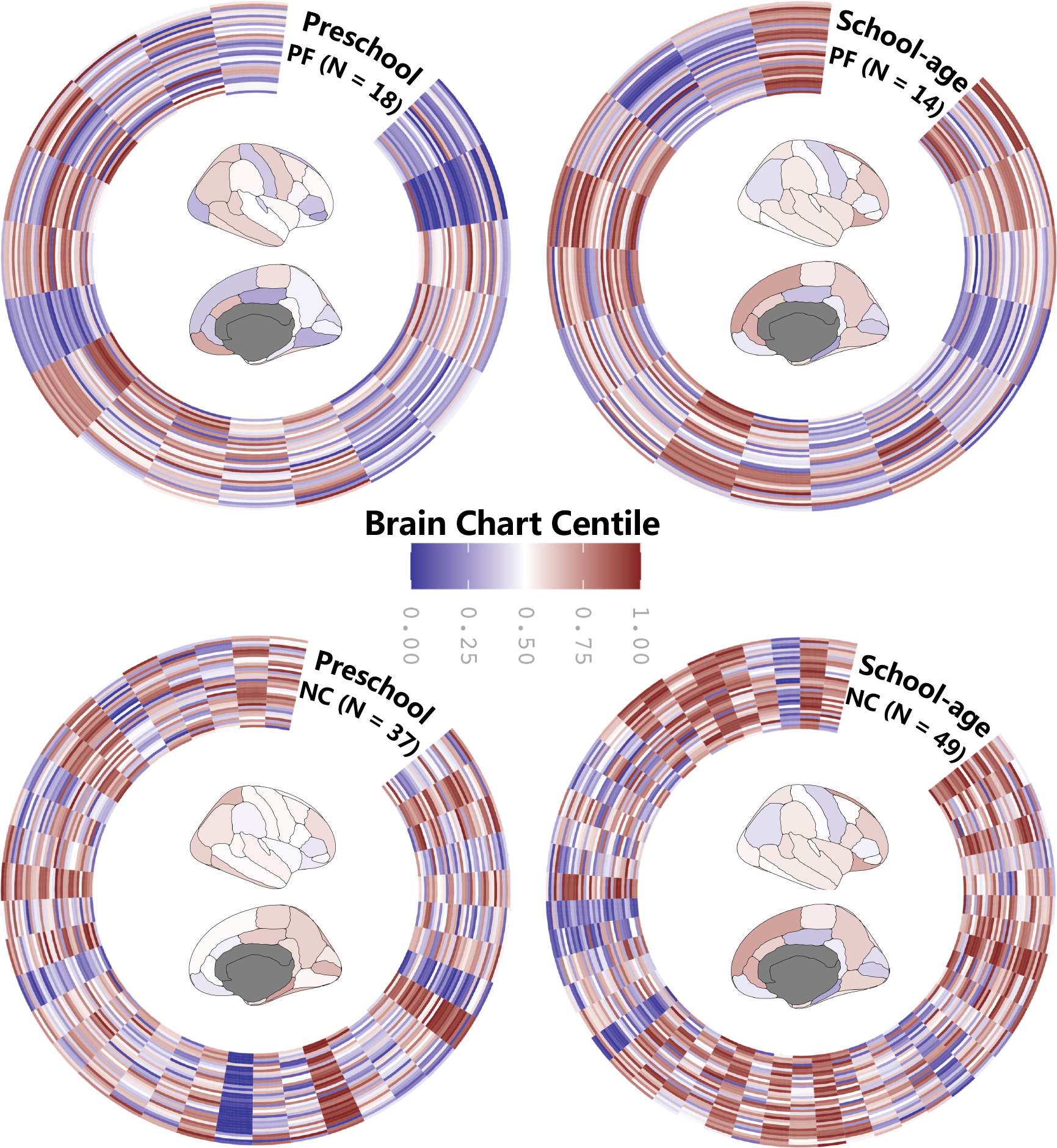
Circular scatter plots of centil scores. The scores of individuals in periCBD were derived from the adjusted lifespan brain charts. Each column in the outer circles represents the centile scores of a region in the Desikan-Killiany (DK) atlas of a child from each of the four (PF preschool, PF school-age, NC preschool and NC school-age) groups. The median centile score is shown for the Desikan-Killiany atlas in the centre for each group.

**Figure 6.**
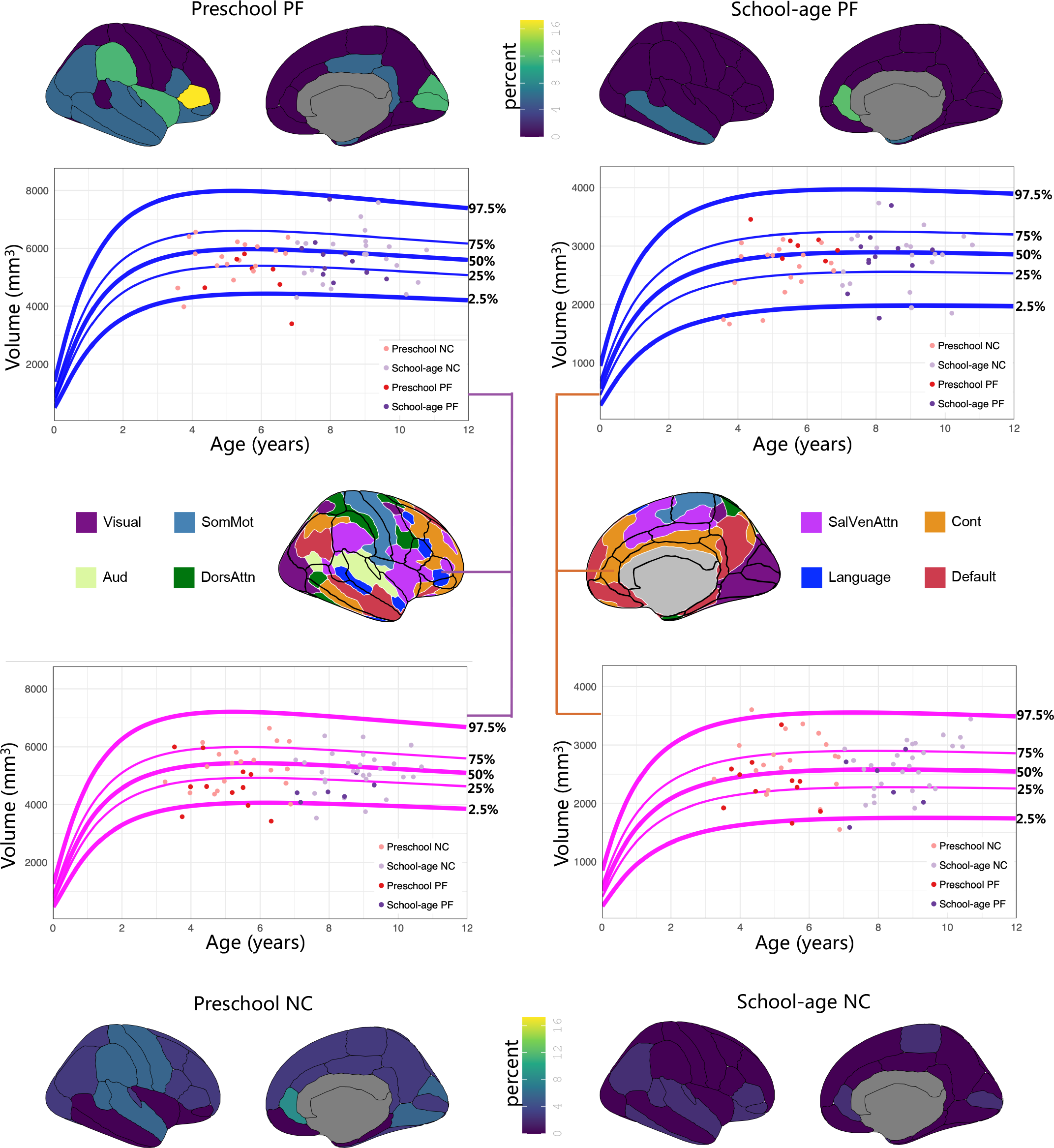
The ratio of regional volume outliers revealed by comparisions with brain growth charts for both the perinatal factor (PF) and normal control (NC) groups during preschool and school age. For each region of the Desikan-Killiany atlas, the percentage of volume outliers was calculated as the ratio of children whose centiles were below 2.5%. Both the parstriangularis of PF preschool children and the rostralanteriorcigulate of PF school-age children (top panel) were outliers given the small size of these regions. Their brain charts are plotted with all the individual centiles rendered as scatter plots for the four groups, with sex indicated by colour (girls =pink charts and boys = blue charts). As shown in the middle panel, we matched the 34 DK regions to the 8 large-scale functional networks^**58**^ to achieve an intuitive understanding of the brain growth chart differences at the network level.

### Brain Chart Analysis

Growth charts for body height, weight and head circumference are widely used to depict age-related trends and further provide developmental benchmarks for pediatric healthcare. Inspired by the pediatric growth charts, growth charts of brain morphology have been developed^49^ by the Lifespan Brain Chart Consortium (LBCC) (https://github.com/brainchart/lifespan) for the human lifespan. These charts allow out-of-sample data from a new MRI study to be aligned with the normative trajectory by estimating specific offsets for the study. This alignment enables the computation of corresponding centile scores for an out-of-sample study.

In this study, the out-of-sample centile scores for six global neurotypes and 34 parcels neurotypes^49^ according to the Desikan-Killiany (DK) atlas^50^ were calculated by estimating sample-specific statistical offsets (i.e., mean *μ*, variance *σ*, and skewness *?*) from age- and sex-appropriate epochs of the normative brain growth trajectory. The modelling was conducted using the generalized additive models for location, scale and shape (GAMLSS)^51^. The centile scores for all neurotypes are shown in Figure 6. The lowest centile score was observed in the supramarginal region of the NC preschool children(1.4 *×*

10^*−*7^), while the highest centile score (0.9999869) was found in the caudalanteriorcingulate region of the NC preschool children. Furthermore, we explored group differences in the median centile scores according to the DK atlas using a two-way ANOVA, with age group (preschool, school-age) and perinatal factors (PF, NC) as between-group factors. The results demonstrated significant main effects of age group (*p* = 0.00049) and perinatal factors (*p* = 0.000003), and revealed a marginally significant effect of the interaction between age group and perinatal factors on median values (*p* = 0.0768).

In addition to analysing centile scores on the brain chart, we also conducted an examination of centile outliers. For each brain phenotype, a centile score below the 2.5% centile was defined as an outlier. The number of outliers for each phenotype was counted and further divided by the number of individuals in each group to obtain the percentile of outliers. As depicted in Figure 6, the results indicate that the preschool groups demonstrated higher rates of extremely small parcels. Among these, the parstriangularis region had the highest deviation in the PF preschool children (16.67%), while the rostralanteriorcingulate region appeared to be the smallest parcel in the NC group (8.11%) . For school-age children, the deviations were relatively low. However, the PF group also demonstrated some larger parcels that exceeded 5%. The parcel with the highest rate in this group was the rostralanteriorcingulate region (11.76%) (Figure 6) .

Taken together, these results demonstrate that there were fewer extreme outliers as children reached school age in both the PF and the NC group. This suggests a transition towards a more “normal” distribution from preschool to school age. However, it is challenging to explain this trend fully given the cross-sectional nature of the dataset presently available. One possible explanation is the effect of parenting and family-based interventions. Parents who have a child with perinatal factors may tend to pay greater attention to their child’s physical training, nutrient intake, and other aspects of paediatric health care. This may also account for the greater number of parcels with lower volumes detected in early childhood, which gradually returned to normal during the course of their growth^52^. Therefore, it is essential to collect longitudinal data in the next stage of periCBD to further explore the underlying mechanisms.

## Usage Notes

### More Perinatal Factors

In this database, weight at birth and gestational age were utilized as the primary perinatal factors. Several other questionnaires were applied that captured perinatal factors across various dimensions. By incorporating these data, researchers can employ a multifactorial analysis strategy to establish associations between brain development and the presence of perinatal factors.

### Developmental Coordination Disorder

Developmental coordination disorder (DCD) is one of the most common NDDs among school-aged children and is characterized by delays in gross and fine motor development without apparent intellectual or medical causes. It is widely believed that DCD predicts the occurrence of various NDDs in K-12 children. Furthermore, some researchers have proposed that DCD may serve as an early indicator of NDDs, considering that motor difficulties/disorders are commonly observed in all types of NDDs. In this dataset, we used the MABC-2 and the DCDQ-C to detect DCD. Researchers can further explore the relationship between NDDs and DCD by leveraging these data.

### Brain-Wide Association Studies with Perinatal Factors

The effects of perinatal factors on the central nervous system have been widely acknowledged, with documented effects such as brain injury and dysmaturation following preterm birth^53^. These factors result in a high degree of heterogeneity in neurodevelopment due to substantial individual differences. Therefore, to elucidate the relationship between perinatal factors and neurodevelopment, it is important to establish brain-wide associations with perinatal factors at an individual level, referred to as individual BWAS-PF^54^. The advent of normative modelling provides an intuitive and effective approach to implement the individual BWAS-PF. By using growth charts derived from extensive data on brain development in healthy individuals, the relative positions of specific brain regions in terms of structural and functional metrics can be obtained, allowing the identification of deviation from the “normal” range^55^. This modelling approach also has promising applications in family intervention and clinical practice^56^. With advancements in normative model algorithms, these models can now be adjusted using out-of-sample MRI datasets with smaller sample sizes^49,57^. This enables the extraction of information on an individual’s relative position in terms of variables such as age, sex, and perinatal factors. Users can leverage this information to establish associations with perinatal factors and other behavioural metrics.

## Code availability

No custom codes or algorithms were used in the generation or processing of data in this manuscript.

## Acknowledgements

The periCBD team receives funding support from the National Basic Science Data Center “Chinese Data-sharing Warehouse for *In-vivo* Imaging Brain” (NBSDC-DB-15), the Key-Area Research and Development Program of Guangdong Province (2019B030335001), the Start-up Funds for Leading Talents at Beijing Normal University, the China Postdoctoral Science Foundation (2022M710432). Gratitude is also extended to the staff at the Anyang Maternal and Child Health Hospital of Henan Province and the People’s Hospital of Liangping District of Chongqing for their support in data acquisition and management. The periCBD team would like to express their appreciation for the research participants and their families who willingly took part in this project. Additionally, the team acknowledges the valuable contribution of all the research assistants involved in data collection, as well as the numerous expert consultants who provided their insights and expertise in the development of the research protocol.

## Author contributions statement

Conception and Design: Qi.Dong., Y.W., X-N.Z., L.K., Q-H.H.

Planning and Discussion: X-N.Z., R-W.H., L.K., Q-H.H., F-M.C., J-H.X., C.Z., S-D.Z., S-Y.Z.

Implementation and Logistics: H-P.A., R.G., S-P.Y., Y.Y., J.L., X-Q.C., X.W., Y-Z.R.

Data Collection: S-P.Y., W.D., S-J.Y., Y.Zhao., Y.Zhou., X-R.F., C-Y.L., Z-Y.W., Y-C.M., W-C.X., L-X.X., X.W., L-F.Z., Z-J.H., H-Y.H., H.L., J.W., Qing.Dong., L-Y.C., S-M,L

Data Informatics: Y-S.W., W.D., S-C.J., S-J.Y., Y-K.Y., P-G., C-Y.L., Z-Y.W.

Quality Control: L-S.L., N-J.J

Data Analysis: Y-S.W., D.C., W.D., S-C.J., L-Z.C., S-J.Y., Y-K.Y., C-Y.L., Z-Y.W., Y-Y.Z., X.Q., F.Z.

Initial Drafting of the Manuscript: Y-S.W., X-T.S. Supervision and Cohort Funding: X-N.Z., L.K.

Critical Review and Editing of the Manuscript: All authors contributed to the critical review and editing of the manuscript.

